# Conflict-mediated tasks for studying the act of facing threats in pursuit of rewards

**DOI:** 10.1101/2020.12.27.424515

**Authors:** Elizabeth Illescas-Huerta, Leticia Ramirez-Lugo, Rodrigo Ordonez Sierra, Jorge A. Quillfeldt, Francisco Sotres-Bayon

**Author notes:** **Correspondence:** Francisco Sotres-Bayon.

## Abstract

Survival depends on the ability of animals to avoid threats and approach rewards. Traditionally, these two opposing motivational systems have been studied separately. In nature, however, they regularly occur simultaneously. When threat- and reward-eliciting stimuli (learned or unlearned) co-occur, a motivational conflict emerges that challenges individuals to weigh available options and execute a single behavioral response (avoid or approach). Previous animal models using approach-avoidance conflicts have often focused on the ability to avoid threats by forgoing the opportunity to obtain rewards. In contrast, behavioral tasks designed to capitalize on the ability to actively choose to execute approach behaviors despite threats are lacking. Thus, we developed three conflict-mediated tasks to directly study rats confronting threats to obtain rewards guided by innate and conditioned cues. One conflict task involves crossing a potentially electrified grid to obtain food on the opposite end of a straight alley, the second task is based on the step-down threat avoidance paradigm, and the third one is a modified version of the open field test. We used diazepam to pharmacologically validate conflict-mediated behaviors in our tasks. We found that, regardless of whether competing stimuli were conditioned or innate, a low diazepam dose facilitated taking action to obtain rewards in the face of threats during conflict, without affecting choice behavior when there was no conflict involved. Using this validated set of innate/learned conflict-mediated tasks could help understand the underlying brain mechanisms that allow animals to confront threats, by actively suppressing defensive responses, to achieve goals.

## Introduction

To ensure survival in nature, animals must avoid threats and pursuit rewards. This ability involves that animals use inherited or assigned value information of stimuli in the environment (negative or positive valence) to control motivated behaviors (Rangel, Camerer et al. 2008, Tye 2018). Traditionally these two motivational valence systems have been successfully studied separately (Hu 2016). Defensive and avoidance responses triggered by threats (LeDoux 2000), have been generally studied separately from approach behaviors elicited by rewards (Cardinal, Parkinson et al. 2002). During foraging, however, animals regularly encounter threats and rewards simultaneously and are consequently challenged to engage in opposing binary choices (avoid or approach) (Choi and Kim 2010, Hayden and Walton 2014, Amir, Lee et al. 2015, Mobbs, Trimmer et al. 2018). Such motivational conflict involves a cost-benefit decision determined by the competition processes between these two mutually exclusive systems interacting. During conflict, individuals must either choose to avoid threats at the cost of not benefiting from rewards or to approach rewards at the cost of facing threats. Traditionally, approach-avoidance conflict tasks have been useful in validating antianxiety drugs like diazepam (Vogel, Beer et al. 1971, Rodgers, Cao et al. 1997, Calhoon and Tye 2015). More recent conflict-mediated animal models have often focused on studying the decision animals make to avoid threats while forgoing the opportunity to obtain rewards (Moscarello and LeDoux 2013, Bravo-Rivera, Roman-Ortiz et al. 2014, Friedman, Homma et al. 2015, Burgos-Robles, Kimchi et al. 2017, Piantadosi, Yeates et al. 2017, Schumacher, Villaruel et al. 2018, Choi, Jean-Richard-Dit-Bressel et al. 2019, Miller, Marcotulli et al. 2019, Verharen, van den Heuvel et al. 2019, Walters, Jubran et al. 2019). Intriguingly, the ability of animals to choose to forage for resources by overcoming threats remains understudied and lack behavioral tasks to characterize it.

To directly study animals seeking rewards in the face of threats, we developed three conflict-mediated choice tasks in rodents based on traditional behavioral assays. All of our behavioral tasks involve comparison of conflict-based versus no-conflict-based choice behaviors. They differ, however, in that rats must use stimuli with innate and/or acquired valences to guide conflict-mediated choices. The crossing-mediated conflict task, based on the task used to map self-stimulation brain sites (Olds and Milner 1954), uses competing cued-conditioned stimuli and involves comparatively assessing the time it takes rats to cross a potentially electrified grid (“threat” zone) to obtain food on the opposite side of a straight alley (“safe” zone) during conflict and no-conflict trials (as mentioned in: Bravo-Rivera and Sotres-Bayon 2020). The step down-mediated conflict task, based on the stepdown threat avoidance paradigm, uses a conditioned aversive event and an innate appetitive stimulus. Finally, the open field-mediated conflict task, based on the open field test (Walsh and Cummins 1976), involves placing food in a brightly lit arena center. We tested the validity of our conflict-mediated tasks by administering systemic injections of a commonly used anticonflict drug, diazepam (Liljequist and Engel 1984). We found that, independently of whether competing stimuli were conditioned or innate, a low dose of diazepam facilitated the ability of rats to overcome a threat incentivized by reward availability during conflict choice behaviors only.

## Methods

All procedures were approved by the Institutional Animal Care and Use Committee of the Universidad Nacional Autónoma de México, in compliance with the National Ministry of Health guidelines for the care of laboratory animals. A total of 153 adult male Wistar rats (Instituto de Fisiología Celular breeding colony) with 2-3 months of age, weighing 250-300 g were housed in individual polyethylene cages, handled daily to diminish stress responses, and maintained on a standard 12 h light/dark schedule. All experiments were performed during the light phase. To maintain a stable motivation to eat or drink during behavioral training and tests, rats were food-restricted (12 g per day of standard laboratory rat chow with a 5 g bonus feeding at the end of each week to maintain rats at 85% of their initial weight) or water-restricted (12 ml per day; 6 ml in the morning and 6 ml in the afternoon), respectively.

### Crossing-Mediated Conflict task

Rats were trained and tested in straight alleys that consisted of acrylic walls with stainless-steel frames (100 cm long x 30 cm wide x 50 cm tall), located in a sound-attenuating cubicle (150 cm long x 70 cm wide x 140 cm tall). The alleys consisted of three zones: two “safe” zones and one “threat” zone (**Fig. 1A**). Each safe zone (20 cm long x 30 cm wide) were located on the opposite ends of the alley and the “threat” zone (60 cm long x 30 cm wide) was located between the two “safe” zones. The floor of the “threat” zone was made of stainless-steel bars (4.8 mm diameter) which delivered a scramble footshock (Coulbourn Instruments, USA), while the floor of “safe” zones was made of acrylic. The “safe” zone included a speaker and standard operant chamber components (cue light, a lever connected to a pellet dispenser and a food plate). The alleys were interfaced with a computer running custom scripts (MATLAB) which controlled apparatus hardware (food pellet delivery, cue lights, speakers and shocker) and recorded task events (lever-presses at each end of the alley). The shock grids as well as the alley floors and walls were cleaned with soapy water, 70% alcohol and dried with paper towels between experiments. Prior to conflict training, all rats were trained to press a lever to obtain sucrose pellets (45 mg, dustless precision pellets, Bioserve, USA) in a fixed reinforcement schedule (each lever press was reinforced with one pellet). All sessions started and ended with context-alone exposure (5 min without cue lights, shocks or noise).

**Figure 1.**
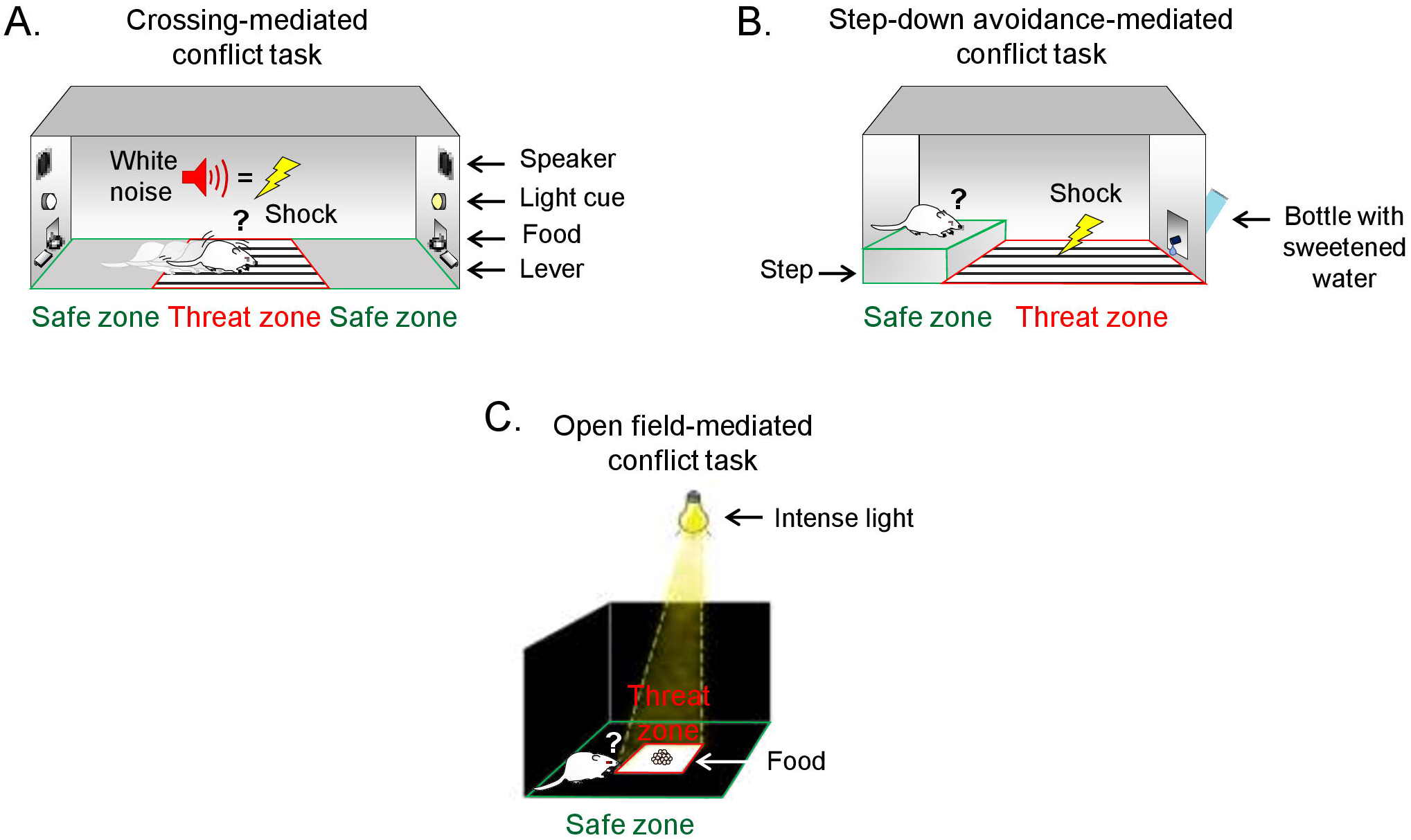
Conflict-mediated tasks to study the ability to confront threats in pursuit of rewards. ***A***, During conflict-mediated crossing task, hungry rats are trained to repeatedly choose to cross a grid (threat zone, red) to press a lever to obtain food (pellets, reward) in the opposite side of the straight alley (safe zone, green), guided by conditioned cues (white noise and light). ***B***, During step-down avoidance conflict task, thirsty rats are trained to choose to step-down from an elevated platform (safe zone, green) onto a conditioned grid (threat zone, red) to approach a bottle containing sweetened water (saccharine, reward). ***C***, During open field-mediated conflict task, hungry rats are challenged to choose to move from the periphery of the arena (safe zone, green) to enter the intensely illuminated center of the arena (threat zone, red) to obtain food (pellets, reward).

### Conflict training

Conflict training involved five succeeding stages: two stages of reward training (reward association and reward crossing), two stages of threat training (threat association and threat crossing), and a final stage of discrimination training to distinguish between conflict and noconflict trials. Conflict training occurred over a total of 30 days.

#### Reward training

Initially, rats confined to one end of the alley (“safe” zone) using an opaque acrylic wall were trained to associate food availability with a light cue (*Reward association*). A pellet was dispensed to the food plate with each lever press when the cue light was illuminated (Light trial), while no pellet was delivered in the absence of light (No-light trial). Each Light trial ended after a randomly assigned number of rewarded pressing events were achieved (ranging from 5-20 presses; a custom script running in a computer associated with the apparatus generated a random number and assigned the type of trial to present), whereas each No-light trial ended after a randomly assigned time had elapsed (ranging from 30 −180 s). Each reward conditioning session was limited to 30 min and it involved ~10 Light trials and ~10 No-light trials which resulted in ~15 min per trial type. One session was given per day across 6 days (days 1 to 6 of conflict training). Next, rats were trained to cross from one end of the alley to the opposite side (from one “safe” zone to the other) to obtain food signaled by light (*Reward crossings*). Short acrylic barriers (9 cm tall) were placed between the grid and “safe” zones to delimit and commit choice-mediated crossing behavior in time and space (choose-point). A reward crossing trial started when a light was illuminated in the side of the alley opposite to the location of the rat. The trial ended when the light was turned off, after either the rat had crossed to the opposite “safe” zone and pressed the lever, or 180 s had passed without crossing. To promote goal-oriented crosses guided by tracking location of light cue, rather than non-signaled habitual reactions, rats received one to three reinforced lever-presses on the same “safe” zone. Each reward crossing session was composed of 30 trials. One session was given per day for 5 days (days 7 to 11 of conflict training). Reward training occurred over a total of 11 days (6 days of reward association and 5 days of reward crossing training). By this point, rats had learned that reward crossings involved actively tracking food availability signaled by the cue light in each of the “safe” zones of the alley in the absence of conflict (crossing to obtain food without threat).

#### Threat training

Rats confined to the “threat” zone (middle of the alley) using opaque acrylic walls and a ceiling (similar dimensions to standard classical threat conditioning chambers: 30 cm long, 25 cm wide, 30 cm tall), were trained to associate a white nose (85 dB for 30 s) with a mild footshock (0.5 mA for 1s) (*Threat association*). To allow the sound coming from opposite sides of the alley to enter the confined conditioning area, the acrylic ceiling was perforated (1.2 cm diameter). Foot-shocks were randomly delivered during the 30-second noise presentation to avoid a temporal noise/shock association that may limit threat-related responses to a specific time (a custom script running in a computer associated with the apparatus generated a random number and delivered the shock). Five noise-shocks pairings were delivered per day with a variable inter-trial interval of 1-3 min. One session was given per day for 4 days. On the next day, two noise-alone trials (in the absence of shocks) were delivered to test for cued threat memory (days 12 to 16 of conflict training). Next, rats were trained to cross the alley to obtain food (guided by the light cue) while the threat cue (noise) was presented simultaneously (*Threat crossings*). A threat crossing trial started and ended the same way as a reward crossing trials. The difference between reward and threat crossings was that during threat crossings rats were challenged to cross to the other side of the alley to obtain food (signaled by conditioned light) by facing the cooccurring threat (0.5mA shock signaled by conditioned noise). Short acrylic barriers (introduced since Reward crossing training) prevented rats from getting exposed to the electrified grid before committing to cross (hesitation at the choice-point). The duration of threat crossing trials was increased across days to gradually intensify threat in crossing sessions. Inter-trial intervals ranged from 1-3 min and each session was limited to 30 min. One session was given per day for 5 days (day 17 to 21 of conflict training). Threat crossing trials lasted 30 seconds in the first two days, 60 s in the third day, 90 s in the fourth day and 120 s in the fifth day of training. Each trial ended either after the time of crossing elapsed (30-120 s) or when rats successfully crossed and pressed the lever on the opposite side of the alley. Threat training occurred over a total of 10 days (5 days of threat association and 5 days of threat crossing training). By this point, rats had learned that threat crossings guided by the simultaneous occurrence of light and sound cues involved conflict (crossing to obtain food despite threat).

#### Discrimination training

The final stage of conflict training consisted of learning to discriminate between No-conflict (crossing to obtain food without threat) and Conflict (crossing to obtain food despite threat) trials across 9 days (*Discrimination*). Each discrimination session consisted on a total of thirty crossing trials. Each session was divided in three blocks of ten trials. Each block of ten trials consisted of seven or nine Noconflict trials and one or three Conflict trials (10% or 30% risk trials, respectively). To prevent rats from predicting types of trials, they were presented in a different sequence across ten-trial blocks (a custom script running in a computer associated with the apparatus generated a random number and assigned the type of trial to present considering the constricted number of types of trials). On each trial, rats were allowed a limited time to choose whether to cross or not to cross the alley (180 s). One session was given per day for 9 days (day 22 to 30 of conflict training). To gradually and progressively increase the risk of threat crossing, proportion conflict trials and intensity of shock increased across days. Conflict crossings were presented with a 10% risk (3 Conflicts versus 27 No-conflict trials) and 0.5mA shock intensity the first three days (22-24 d), the next three days (25-27 d) with a 30% risk (9 Conflict versus 21 No-conflict trials) using the same shock intensity as the sessions before, and finally the last three days (28-30 d) with the same percentage of risk as the session before but increased intensity of shock of 0.1mA per day until it reached 0.8mA at the end of discrimination training. At this point, rats had learned to discriminate between Conflict against No-conflict trials. Rats that did not learn to discriminate between types of trials by the end of discrimination training (p > 0.05 in the comparison between Conflict and No-conflict trials in the last block of trials) were excluded from the study (4 out of 18).

### Conflict test

The conflict test was performed one day after conflict training ended. Rats were tested with 10 crossing trials. These trials were presented in the same conditions as the last day of discrimination training (30% risk: 3 conflict and 7 no-conflict pseudorandomly presented trials) but in the absence of shocks. Results were expressed comparing the average of Conflict against No-conflict trials before (pre-Test) and after (Test) injections.

### Step-down avoidance-mediated conflict task

Rats were trained and tested in a modified step-down inhibitory avoidance chamber (50 cm long x 25 cm wide x 25 cm tall) located in a sound-attenuating cubicle (61.5 cm long x 62.5 cm wide x 65 cm tall). The step-down chamber consisted of two zones: one “safe” and one “threat” zone (**Fig. 1B**). The floor of the “threat” zone (30 cm long x 25 cm wide) consisted of stainless-steel bars (4.8 mm diameter) delivering a scrambled footshock (Coulbourn Instruments, USA), while the floor of the “safe” zone (20 cm long x 23.5 cm wide x 6 cm tall) was an acrylic-covered elevated step platform. An acrylic sliding door was located between the threat and safe zones to allow precise timing of step-down choice behavior (choice-point). Compared to the standard step-down avoidance (Izquierdo, Quillfeldt et al. 1997), the modified chamber includes a bottle with 12 ml of saccharin solution (0.1%) placed on the wall of the “threat” zone opposite from the “safe” zone. This modification represents an incentive that motivates the animal to step-down from the safe platform.

Before training started, rats were habituated to the step-down behavior in the chamber for one day (the bottle was present but empty) and water-deprived for 12 hours. Training and test sessions started with rats confined to the elevated platform for 5 min on the safe “zone” (sliding door was closed). Shock grids as well as chamber floors and walls were cleaned with soapy water, 70% alcohol and dried with paper towels between experiments.

### Conflict training

Two types of step-down tasks were used in separate groups of rats. One task involved conflict training, while the other task did not involve conflict. The only difference between these tasks is that conflict training involved water-restricted rats (thirsty), while rats in the no-conflict task had free access to water (satiated). Both tasks involved reward presentations and threat association training. During reward presentation sessions, after pulling the sliding door, rats were allowed to drink from the bottle placed in the opposite side of the elevated platform (“threat” zone) containing saccharin solution (12 ml). Each reward presentation session was limited to 10 min. One session was given per day for 5 days (days 1 to 5), to allow familiarization with the chamber, overcoming neophobia to the novel taste (saccharine) and achieving stable reward sampling across days. The following two days (days 6 and 7), rats were trained to associate the act of stepping down from the platform with a mild footshock (0.5 mA). This footshock lasted until rats came back to the safety platform. One trial per session was given per day. Step-down training for each of the two tasks (conflict and no-conflict) occurred over a total of 7 days. By this point, two separate groups of rats had been trained in two different conditions: one group (motivated to drink) had learned to step-down to obtain sweetened water despite threat (conflict group) while another group (not motivated to drink) had learned to avoid threat by not stepping down (noconflict group).

### Conflict test

A test that involved conflict and another that did not involve conflict were performed one day after training ended. Rats were tested in the same conditions of the last day of training of each of the groups (Conflict and No-conflict) but in the absence of shocks. Results were expressed comparing the average of Conflict and No-conflict group step-down latencies before (pre-Test) and after (Test) injections.

### Open Field-Mediated Conflict task

Rats were tested in a modified open field arena (90 cm long, 90 cm wide, 60 cm tall) with walls made of wood and floors made of textured transparent acrylic, located in a dark room. The arena was divided into two zones: one “safe” zone (60 cm long, 60 cm wide) and one “threat” zone (30 cm long, 30 cm wide) (**Fig. 1C**). Compared to the standard open field (Stefanski, Palejko et al. 1992) our modified arena includes an intense beam of light (1500 lux) focused to the center of the arena (“threat” zone), allowing the periphery of the arena to remain dark (“safe” zone). This modification represents an added innately aversive stimulus to the already innately aversive center of the open field without affecting the periphery of the arena. The floor and wooden walls were cleaned with soapy water, 70% alcohol and dried with paper towels between experiments.

### Conflict test

Two types of open field tests were used in separate groups of rats. One test involved conflict, while the other test did not involve conflict. The only difference between these two tests is that the Conflict test involved placing thirty sucrose pellets (1.350 g) in the center of the field (“threat” zone) to motivate rats to forage for food (Britton and Britton 1981), whereas the No-conflict test did not involve food in the arena. Motivation to forage for food was maintained in the periphery by placing five sucrose pellets (0.9 g) on each of the corners of the arena. To avoid innate aversion to the novel taste (neophobia) of pellets, rats received twenty sucrose pellets (1.125 g) in their home cages for two days before tests. Rats were individually placed into the “safe” zone, and their time spent in the “threat” and the “safe” zones was recorded for five minutes. The time spent in the center of the open field arena was used to evaluate foraging despite threat (Conflict) and time spent in the periphery was used to evaluate foraging without threat (No-conflict). Test results were expressed comparing the average times spent on either center or periphery of the arena after injections in conflict and no-conflict groups.

#### Open field test

Locomotor activity and anxiety were evaluated in a standard open field arena (no food or intense light involved)(Schmitt and Hiemke 1998). Rats were individually placed in the center of the open field arena, and their behavior was recorded for 5 min. Distance travelled in the open field was used to evaluate locomotion, and entries in the center of the arena were used to evaluate anxiety-like behavior.

#### Beam walking test

Motor coordination was evaluated using beam walking behavior (Goldstein and Davis 1990). The beam was a wooden bar (100 cm long x 2 cm wide) placed 80 cm above the floor with an inclination of 15°. An opened home cage was placed at the end of the beam to motivate rats to walk. The latencies to arrive at the end of the beam and enter into the home cage were recorded. Rats were trained for five days to walk on the beam towards the home cage, initially (day 1 and 2) starting at a distance of 50 cm and then (day 3 to 5) starting at a distance of 100 cm from the home cage. Rats were allowed to stay in the home cage for 30 seconds. The next day, rats were tested on beam walking.

#### Food intake test

Feeding behavior was evaluated by calculating food intake in rats single housed in their home cages (Britton and Britton 1981, Rex, Stephens et al. 1996). Food consumption was assessed across three days. Each day, rats were presented with 30 gr of sucrose pellets placed in a familiar food-plate. To calculate food intake, the food plate was weighed before and after 30 min of food presentation.

### Systemic drug administration

A subcutaneous dose (1 or 2 mg/kg of weight) of diazepam (DZPM, Valium ^®^, Roche; México) or saline solution (SAL) was administered 30 min before all behavioral tests.

### Data collection and analysis

All behavior was recorded with digital video cameras (Provision, model D-380D5) located above each of the task apparatus. Lever pressing events were recorded by a computer running MATLAB^®^ (MathWorks Inc.) custom scripts interfaced with the straight alley. Lever pressing events (presses per minute) during light or no-light trials were expressed as blocks of ten minutes. Each session (one per day) consisted of three of these ten-minute blocks.

Crossing latencies consisted of the total time (seconds) that rats spent to cross to the opposite side of the straight alley and press the lever to obtain food. Latencies to cross were expressed as blocks of ten trials. Each training session (one per day) consisted of three of these ten-trial blocks and the test session consisted of one block of ten trials. Freezing responses were expressed as the percent of time spent without movement (except for respiration) during context alone exposure (5 min baseline before noise presentations) or sixty seconds after noise presentations. Step-down latencies were expressed as the total time (seconds) rats took, after pulling the sliding door, to step-down from the elevated platform with four paws onto the grid. Beam walking latencies consisted of the total time (seconds) spent before arriving at the end of the beam to enter into the home cage. Food intake was calculated by comparing food-plate weight (grams) before and after thirty minutes of food presentation. Time spent freezing, as well as step-down, beam walking latencies and food intake were manually recorded by trained observers blinded to experimental conditions. Behavior in the open field arenas was recorded using a tracking software (ANY-maze; Stoelting, USA) and expressed as travelled distance (cm) or time spent (seconds) in the center or periphery of the field during 5 min. After testing for the normality of data (Kolmogorov-Smirnov test), experimental groups were compared by using, when appropriate, Student’s two-tailed *t* tests (paired or unpaired) or analysis of variance (one- or two-way ANOVA) followed by planned comparisons or post hoc Tukey’s honest significant difference test (STATISTICA; StatSoft, USA).

## Results

To determine which dose of diazepam was appropriate to test the validity of our conflict-mediated tasks, we started by evaluating two diazepam doses in two commonly used behavioral assays. We injected rats with diazepam (DZPM) at a low dose (1mg/kg), a high dose (2mg/kg) or saline solution (SAL) before open field (low dose, *n* = 8; high dose, *n* = 9; SAL, *n* = 17) and beam walking tests (low dose, *n* = 6; high dose, *n* = 8; SAL, *n* = 6). We found that diazepam at a low dose did not affect locomotion, anxiety-like behavior nor motor coordination, as indicated, respectively, by similar levels of total distance traveled in the periphery (SAL: 212.8 cm, DZPM: 217.4cm; *t*_(14)_ = −0.11, *p* = 0.910) and number of entries to the center of the open field (SAL: 1.6, DZPM: 1.0; *t*_(14)_ = 0.70, *p* = 0.491), as well as similar times to arrive to the end of a narrow elevated beam, as compared to the control group (SAL: 9.6 s, DZPM: 12.8 s; *t*_(10)_ = −2.04, *p* = 0.068). In contrast, rats injected with high diazepam dose impaired locomotion and motor coordination, without affecting anxiety-like behavior, as indicated by decreased levels of total distance traveled in the periphery (SAL: 231.8 cm; DZPM: 103.6 cm; *t*_(16)_ = 3.07, *p* = 0.010) and similar number of entries to the center of the open field (SAL: 2.3; DZPM: 1.1; *t*_(16)_ = 1.42, *p* = 0.173), as well as decreased times to arrive to the end of a narrow elevated beam (SAL: 9.66 s; DZPM: 120.7 s; *t*_(12)_ = − 3.28, *p* = 0.006), as compared to control group. Given the impairment effects on locomotion and motor coordination observed after injecting a high dose of diazepam, we used a low dose of diazepam to test the validity of conflict-mediated behaviors in our tasks.

### Diazepam facilitated crossing during conflict test without affecting no-conflict behaviors

To evaluate the effect of diazepam on conflict-mediated crossing behavior at test, rats received conflict training in three consecutive stages: reward and threat trainings, followed by trial discrimination training (**Fig. 2A**). Threat and reward trainings involved associative and crossing sessions. During reward association sessions (6 days), rats confined to one of the ends of the straight alley (“safe” zones), gradually learned that a light cue was associated with availability of food after pressing a lever, whereas the absence of light was associated with lack of food availability, as indicated by the progressively decreasing lever pressing levels in No-light trials as compared to stable lever pressing in Light trials across training days (group: *F*_(1,26)_ = 12.3, *p* = 0.001; trial block: *F*_(17,442)_ = 6.09, *p* < 0.001; interaction: *F*_(17,442)_ = 24.0, *p* < 0.001). Notice that reward conditioning shifted an apparent natural preference for lever pressing during the absence of light cue at the beginning of training (day 1, first trial block: Light: 7.58, no-Light: 11.63 presses/min; *t*_(13)_ = −2.69, *p* = 0.018), to stable pressing preference guided by the light cue at the end of training compared to trials in the absence of light (day 6, session average: Light:10.12, no-Light: 4.49 presses/min; *t*_(13)_ = 7.79, *p* < 0.001). During reward crossing sessions (5 days), the alley was opened to allow for rats to learn to track food availability on opposite ends of the alley guided by the light cue, as indicated by a sustained decrease of time spent to choose to cross the alley (latency) across days (*F*_(14,182)_ = 16.4, *p* < 0.001). Notice that rats rapidly learned to cross for food cued by light, as indicated by long latencies in the beginning of training (day 7, first trial block: 20.4 s) as compared to short stable latencies by the end of training (day 11, session average: 8.7 s; *t*_(13)_ = 6.11, *p* < 0.001).

**Figure 2.**
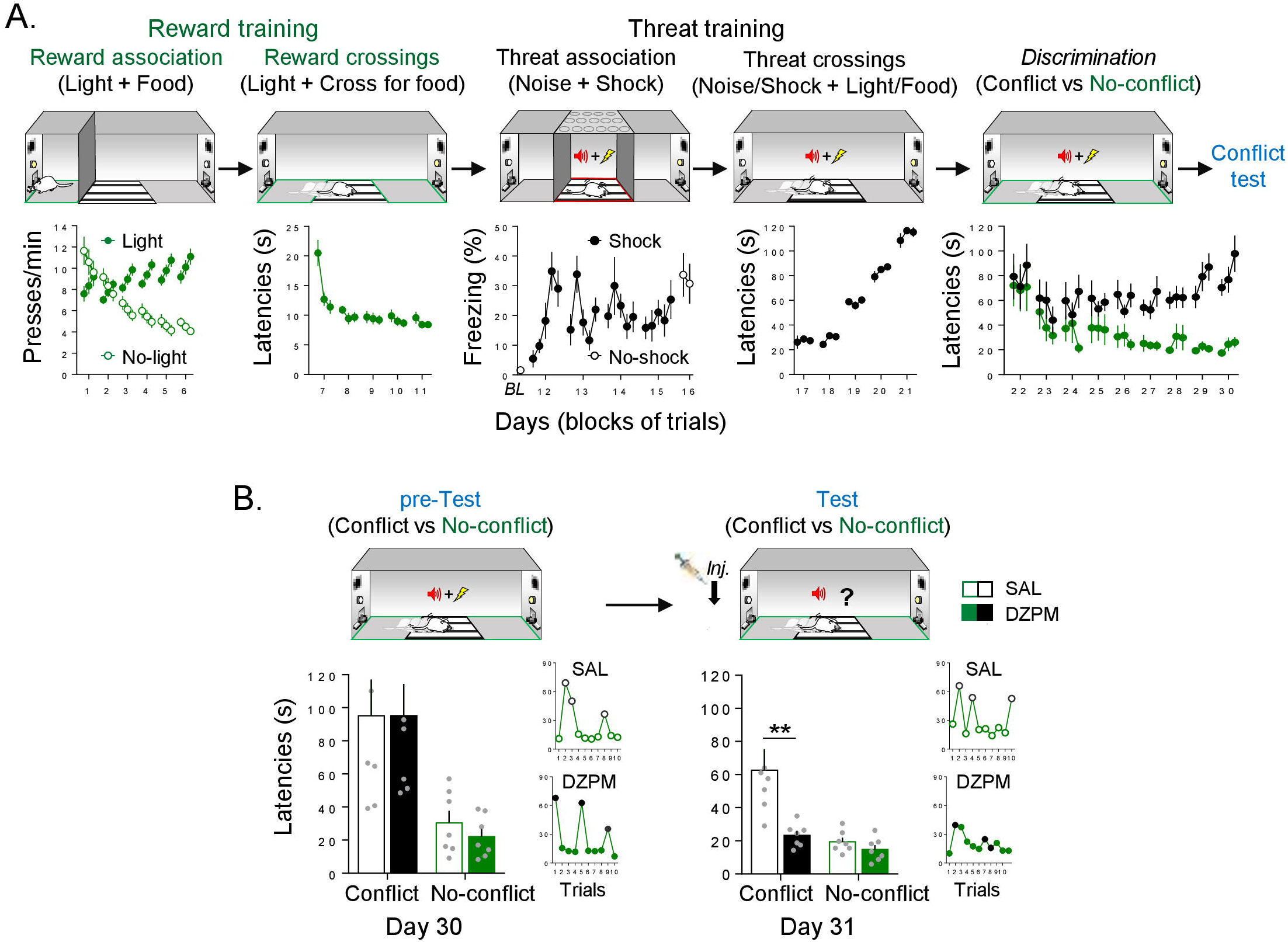
Diazepam decreases crossing latencies during conflict test without affecting noconflict trials. ***A***, Rats acquired conflict-mediated crossing in 30 days (n=14). First, hungry rats, confined to the safe zone (green), learned to associate pressing a lever with food availability cued by a light (reward conditioning), followed by training to cross to the opposite safe zone of the straight alley to obtain food cued by light (no-conflict crossings). Then, rats, confined to the threat zone (grid, red) of the alley, learned to associate occurrence of white noise with a mild footshock (threat conditioning), followed by training to cross with both learned contingencies (light/food and noise/shock) presented simultaneously (conflict crossings). Finally, rats were trained to discriminate crossing trials guided by no-conflict (light/food alone) or conflict (light/food and noise/shock) cues. Data from lever pressing (per minute) and time to cross to the opposite safe zone of the alley (latency in seconds) are presented in blocks of three trials per day, whereas percent time spent freezing (with or without shock) is presented for each trial. By the end of conflict-mediated crossing training, rats showed high crossing latencies during conflict trials (black) compared to no-conflict trials (green). ***B***, Before injection (pre-Test), saline solution and diazepam groups (SAL, n = 7; DZPM, n = 7) showed similarly high crossing latencies during conflict trials and similarly low latencies during no-conflict (*Left*, trials averages; *Right* trial by trial performance of representative rats). The following day, after injection (Test), diazepam-treated rats decreased crossing latencies during conflict trials (no shock), while leaving no-conflict trials intact, as compared to saline-treated rats (*Left*, trial averages; *Right*, trial by trial performance of the same representative rats shown in pre-test). Error bars indicated SEM. BL, baseline. ** *p* < 0.01.

Threat training initiated one day after reward training. During threat association sessions (5 days), rats, confined to the middle of the alley (“threat” zone), were conditioned and tested. Rats rapidly learned to associate noise presentations with shock occurrence, as indicated by increased freezing levels across the first day of conditioning (day 12, *t*_(13)_ = − 5.28 *p* < 0.001). Consistent with this learning, rats showed robust threat memory retrieval elicited by noise in the absence of shock, as indicated by high freezing levels during the test compared to pre-conditioning baseline levels (day 16, *t*_(13)_ = −5.23 *p* < 0.001). Between initial conditioning and test, rats exhibited escape-like behaviors as indicated by variable freezing levels (days 13 to 15). Such threat-elicited behavior reflects the unexpected shock delivery timing during noise presentations (see Methods) intended to prime active, rather than passive, defensive responses when challenged to cross the alley. Starting the next day, the alley was opened to allow crossings. During threat crossing training (5 days), rats learned to cross to the opposite end of the alley to obtain food despite an electrified grid signaled by noise presentations, as indicated by increased crossing latencies across days (*F*_(14,182)_ = 142.4, *p* < 0.001). By gradually and progressively, increasing the duration of time allowed to cross the grid across days (see Methods), rats learned to limit threat crossing response timing in the choice-point, as indicated by short latencies in the beginning (day 17 first trial block: 26.1 s) compared to long latencies at the end of training (day 21 session average: 113.3 s; *t*_(13)_ = −16.6, *p* < 0.001).

These results indicate that, at this point, rats had learned reward and threat contingencies separately. In the final conflict training stage, rats received discrimination training (9 days). Rats learned to distinguish, in the same session, between pseudorandomly presented crossing trials that involved conflict (light and noise presented simultaneously) against those that occurred in the absence of conflict (light without noise), as indicated by increasingly long latencies during Conflict and low latencies during Noconflict trials across days (group: *F*_(1,15)_ = 34.2, *p* < 0.001; trial block: *F*_(26,390)_ = 2.72, *p* < 0.001; interaction: *F*_(26,390)_ = 1.53, *p* = 0.047). Notice that rats started training with similar latencies between trial types (day 22 session trial averages: Conflict, 94.0 s vs No-conflict 78.8 s; *t*_(13)_ = −1.66, *p* = 0.119) and ended (day 30 session average) with prominently high latencies during Conflict trials (81.6 s) as compared to low latencies during No-conflict trials (22.7 s; *t*_(13)_ = −8.00, *p* < 0.001).

To evaluate the effect of diazepam on conflict-mediated crossing behavior at test, rats were separated into two groups matched by similar crossing latencies within trial types and maintained differences between trial types (pre-Test, group: *F*_(1,24)_ = 0.07, *p* = 0.787; trials: *F*_(1,24)_ = 20.65, *p* < 0.001; interaction: *F*_(1,24)_ = 0.076, *p* = 0.784) (**Fig. 2B**). The next day (day 31), rats were tested with ten crossing trials after SAL or DZPM injections. Notably, we found that diazepam decreased crossing latencies during conflict test without affecting noconflict trials, as indicated by low latencies in diazepam-injected rats (23.3 s) compared to long latencies in saline-injected rats (62.6 s) during conflict trials, yet similar latencies between groups during No-conflict trials (DZPM: 14.7 s, SAL: 19.4 s) (Test, group: *F*_(1,24)_ = 10.6, *p* = 0.003; trial block: *F*_(1,24)_ = 14.75; *p* < 0.001; interaction: *F*_(1,24)_ = 6.58; *p* = 0.016). Notice that diazepam-treated rats crossed as rapidly when conflict was involved as when conflict was absent, as indicated by similar latencies in conflict as compared to No-conflict trials during test after diazepam injection (23.3 s and 14.7 s latencies, respectively; *post hoc* comparison after ANOVA: *p* = 0.80). Figure 2B shows trial by trial performance comparison of two individual representative rats before (pre-Test) and after (Test) injection of either SAL or DZPM, highlighting that the systemic drug manipulation facilitated alley crossings that involve selecting the appropriate choice-behavior guided by competing threat and reward memories. To test whether this diazepam effect could be explained by impairment on expression of threat and/or reward memories independently, we injected saline solution or diazepam in separate groups of rats before threat (SAL, *n*=5; DZPM, *n* = 5) and reward (SAL, *n* = 6; DZPM, n = 7) conditioning retrieval tests. Separate groups of rats were subjected to threat conditioning and reward conditioning as above (see Threat association and Reward association training, respectively) and tested for memory retrieval on the last day after injection of either diazepam or saline solution. We found that diazepam, did not affect threat and reward memory expression, as indicate by similar freezing (DZPM: 57.33, SAL: 51.6 percent freezing) and lever pressing levels (DZPM: 8.7, SAL: 9.6 presses/min) compared to their respective saline groups during the retrieval test (*t*_(8)_ = −0.49, *p* = 0.63; *t*_(11)_ = −1.03, *p* = 0.32). Finally, we evaluated whether crossing-mediated choice behavior during the conflict test was goal-directed or habitual by retraining rats for three days, giving rats free access to food for a day and the following day evaluated for crossing behavior. We confirmed that crossing behavior was guided by goals and not habits, as indicated by maintained trial discrimination but increased latencies in both conflict and no-conflict trials in satiated (Conflict trials: 154 s; No-conflict trials: 59 s) versus no-satiated rats (Conflict trials: 64.6s; No-conflict trials: 17 s) during test (comparison: Conflict trials, *t*_(13)_ = −5.43, *p* < 0.001, No-conflict trials, *t*_(13)_ = −684, *p* < 0.001). These results indicate that diazepam does not affect threat- and reward-related behaviors *per se*, but rather facilitates goal-directed conflict-mediated behaviors that involve simultaneously occurring threat and reward predicting stimuli that compete for the selection of the appropriate choice behavior. Thus, taken together, these findings validate our crossing-mediated conflict task by showing that diazepam facilitates the ability of rats to choose to confront learned threats to execute learned motivational responses that lead to obtaining food.

### Diazepam facilitated step-down latencies during conflict test without affecting noconflict behaviors

Facing threats to obtain rewards are often guided, not only by stimuli that acquire valence through learning but, by naturally preferred stimuli. Animals often forage for rewards that possess innate (unlearned) value, such as water and sweet tastes. To study animals seeking an innate reward in face of a learned threat, we evaluated retrieval of an aversive memory in conflict-based versus no-conflict based step-down avoidance tasks. In the standard step-down avoidance task, rats rapidly learn that stepping down from a safe platform that predicts a footshock in an electrified grid floor (Izquierdo, Quillfeldt et al. 1997). To study conflict-mediated behavior, we developed a modified version of the step-down avoidance task that involves placing a bottle containing naturally preferred sweetened water at the end of the grid, opposite to the safe platform (see Methods).

To evaluate conflict-mediated step-down avoidance behavior at test, we compared choice behavior (latencies to step down or not to step down the platform to obtain water with saccharin) in two separate groups of rats. One group of rats received conflict training, while the other group received no-conflict training. Conflict training involves rats that are motivated to face the learned threat because they are thirsty, whereas no-conflict training involves rats that had free access to water (satiated) and therefore were not motivated to step-down the safe platform. Both groups of rats received conflict and no-conflict training in two consecutive stages: reward presentation and threat training (**Fig. 3A**). During reward presentation (5 days), rats gradually learned that at the end of the grid there was a bottle of water containing saccharin, as indicated by the progressively decreasing latencies in both conflict (*F*_(1,64)_ = 17.63; *p* < 0.001) and no-conflict training (*F*_(1,44)_ = 11.35; *p* < 0.001). In a few days, rats familiarized with the box and identified the location of the reward, as indicated by long latencies in the first reward presentation (day 1, conflict training: 284.4 s; day 1, no-conflict training: 463.0 s) as compared to short latencies by the last reward presentation session (day 5, conflict training: 15.5 s; day 5, no-conflict training: 101.0 s; conflict training: *t*_(16)_ = 4.53, *p* < 0.001; no-conflict training: *t*_(11)_ = 5.04, *p* < 0.001). At this point, both groups of rats showed similarly similar step-down latencies to obtain saccharin (*t*_(27)_ = −1.73, *p* = 0.094).

**Figure 3.**
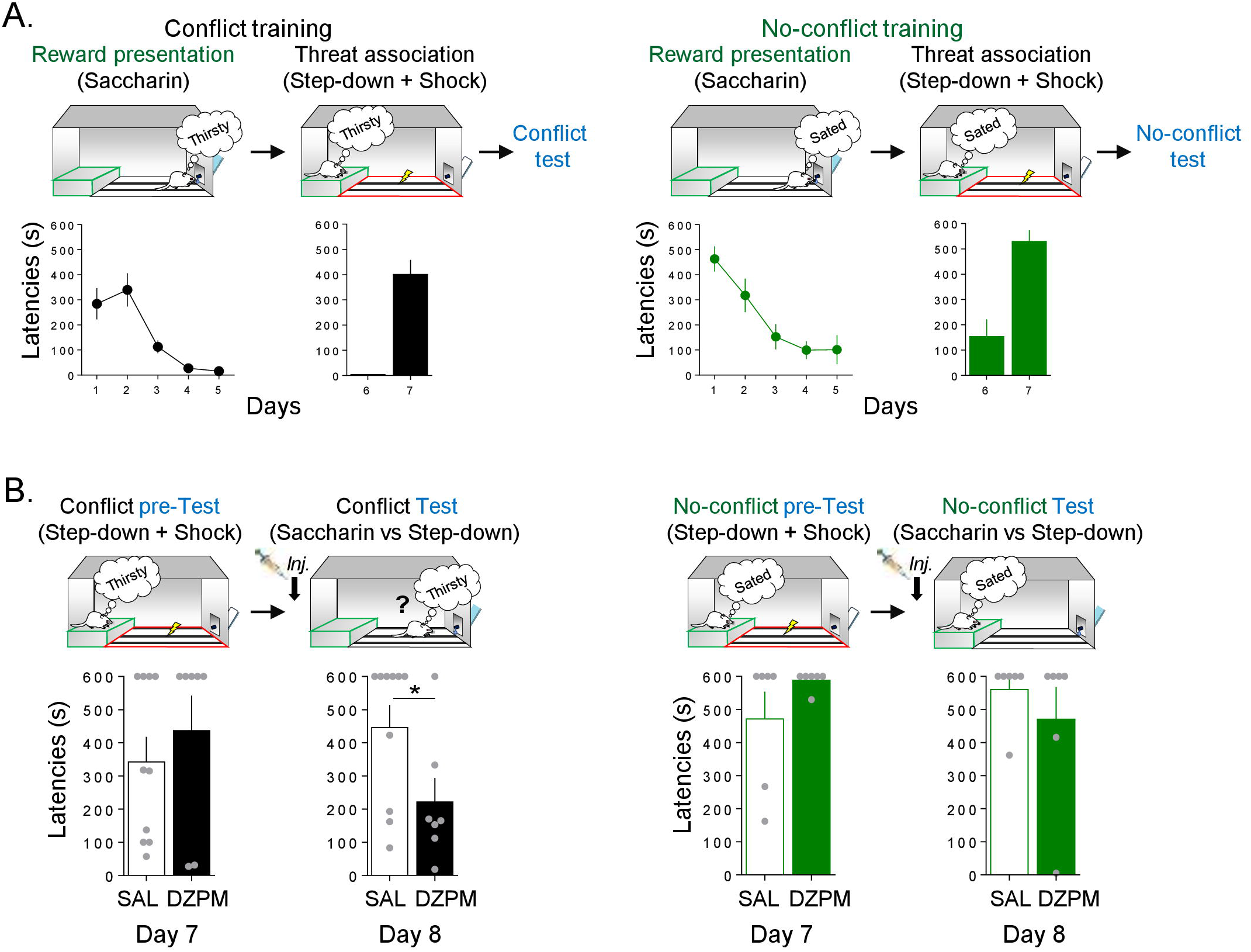
Diazepam decreases step-down latencies during conflict test without affecting the no-conflict condition. ***A***, Rats acquired step-down avoidance either mediated by conflict or no-conflict. *Left*, In the conflict condition (black), rats (n = 17) innately motivated to drink (thirsty) step-down the platform (safe zone, green) to obtain sweetened water (saccharin; reward presentation) from the bottle at the end of the grid, followed by learning that the act of stepping down was associated with the occurrence of a mild footshock in the grid context (threat zone, red). *Right*, In the no-conflict condition (green), rats (n = 12) with free access to water (sated) stepped down the platform to obtain saccharin solution, followed by learning that stepping down was associated with footshock delivery in the grid. Time to stepdown with four paws onto the grid (latency in seconds) is presented by a single trial per day. By the end of training, both groups showed high latencies to step-down. ***B***, Before injection (pre-Test), saline solution (SAL) and diazepam (DZPM) groups, in both conflict (*Left*, black) and no-conflict (*Right*, green) conditions, showed high latencies to step-down. The following day, after injection (Test), DZPM-treated rats decreased step-down latencies during the conflict condition (*n* = 7) without affecting the no-conflict condition (*n* = 6), as compared to SAL-treated rats (*n* = 10 and *n* = 6, respectively). Error bars indicated SEM. * *p*< 0.05.

Threat association was initiated one day after the reward presentation. During threat association sessions (2 days), rats received a mild footshock in the grid (“threat” zone) immediately after they stepped down from the platform (“safe” zone). Rats rapidly learned to associate the action of stepping down from the platform with footshock occurrence, as indicated by short latencies in the beginning (day 6, conflict training: 4.1 s, day 6, no-conflict training: 154. 0 s) compared to long latencies at the end of training in both group of rats (day 7, conflict training: 401.0 s; *t*_(16)_ = −6.97, *p* < 0.001; day 7, no-conflict training: 529.9 s; *t*_(11)_ = −5.64, *p* < 0.001). At this point, rats had associated the step-down response with footshock.

To evaluate the effect of diazepam on conflict-mediated step-down behavior at test, rats that received conflict or no-conflict training were separated into two groups matched by similar step-down latencies (day 7, pre-Test, conflict group: *t*_(15)_ = −0.74, *p* = 0.46; day 7, pre-Test, no-conflict group: *t*_(10)_ = −1.40; *p* = 0.19; **Fig. 3B**). The next day (day 8), rats were tested after SAL or DZPM injections. Notably, we found that diazepam decreased stepdown latencies during conflict test, as indicated by low latencies in diazepam-injected rats compared with saline-injected rats (SAL: 446.1 s, DZPM: 225.5 s; *t*_(15)_ = 2.2; *p* = 0.043). In contrast, diazepam did not affect step-down latencies during no-conflict test, as indicated by similar long latencies in diazepam-injected rats (s) compared with saline group (SAL: 560.3 s, DZPM: 470.1 s; *t*_(10)_ = 0.85; *p* = 0.41). These results indicated that diazepam decreases step-down avoidance responses during conflict without affecting the expression of avoidance memory when there was no conflict involved. Thus, taken together, these findings validate our step-down avoidance-mediated conflict task by showing that diazepam facilitates the ability of rats to choose to overcome a learned defensive response to actively obtain a naturally preferred reward.

### Diazepam facilitated foraging behavior during conflict test without affecting noconflict behaviors

In nature, foraging behavior often requires that animals overcome the risk of encountering a predator. Rodents, for example, may be challenged to forage for food in an open field despite the risk of being detected by a flying predator. To study the ability of rodents to face innate threats to obtain food, we evaluated foraging behavior in conflict-versus no-conflict-based open field tests. In the standard open field test, rats explore the periphery of a novel environment (“safe” zone) while innately avoiding the center of the open field arena (“threat” zone) (Prut and Belzung 2003). To study conflict-mediated behavior, we developed a modified version of the open field test by placing food in a brightly lit center arena (see Methods).

To evaluate conflict-mediated foraging behavior, we compared open field behavior (time spent in periphery and center of an open field arena) in two separate groups of rats. One group of rats was exposed to conflict conditions (Conflict), while the other group was exposed to the same environment in the absence of conflict (No-conflict). Conflict test involves food in the center of the open field arena (**Fig. 4A**), whereas No-conflict test does not (**Fig. 4B**). To evaluate the effect of diazepam on conflict-mediated foraging behavior, we compared open field behavior in conflict-mediated versus no-conflict mediated open field tests after SAL and DZPM injections. We found that diazepam increased foraging behavior for food during conflict test as indicated by more time spent in the center of the open field (“threat” zone) in diazepam-injected rats as compared to saline-injected rats (SAL: 70.85 s, DZPM: 116.9 s; *t*_(19)_ = −3.9, *p* < 0.001). As a result of increased foraging behavior in the center, diazepam-injected rats spent less time in the periphery (“safe” zone) than saline-injected rats (SAL: 229.1 s, DZPM: 183.7 s; *t*_(19)_ = 3.9, *p* < 0.001). In contrast, diazepam did not affect foraging behavior during no-conflict test, as indicated by both diazepam and saline-injected rats spending similar time in the center (SAL: 19.3 s, DZPM: 11.7 s; *t*_(10)_ = 0.96, *p* = 0.35) and periphery (SAL: 280.6 s, DZPM: 288.2 s; *t*_(10)_ = −0.96, *p* = 0.35) of the open field arena. Diazepam effect on conflict test was not due to increased food intake, as indicated by similar levels of food consumed compared to the saline group during the conflict test (SAL: 2.23 g, DZPM: 2.24 g; *t*_(19)_ = 0.68, *p* = 0.500). To further test whether the low diazepam dose effect could be explained by hyperphagia (Johnson 1978, Naruse and Ishii 1995), we injected saline or diazepam in separate groups of rats before food intake test. We found that diazepam did not increase food intake, as indicated by similar levels of food consumed compared to saline-treated group in the third day of food intake test (SAL (*n* = 4): 16.7 g, DZPM (*n* = 4): 17.7 g; *t*_(6)_ = −0.35, *p* = 0.73). Consistently, by comparing food intake in the same individual during drug-free test (day 2) and after diazepam test (day 3), we found that diazepam did not affect food intake, as indicated by similar weight of food-containing plates between days (day 2 (without drug): 15.0 g, day 3 (with drug): 17.7 g; *t*_(3)_ = −1.06, *p*= 0.36). These results indicate that diazepam increases foraging behavior during conflict without affecting no-conflict behaviors (including feeding). Taken together, these findings validate conflict behavior in our open field-mediated conflict task by showing that diazepam facilitates the ability of rats to choose to confront innate threats to forage for food.

**Figure 4.**
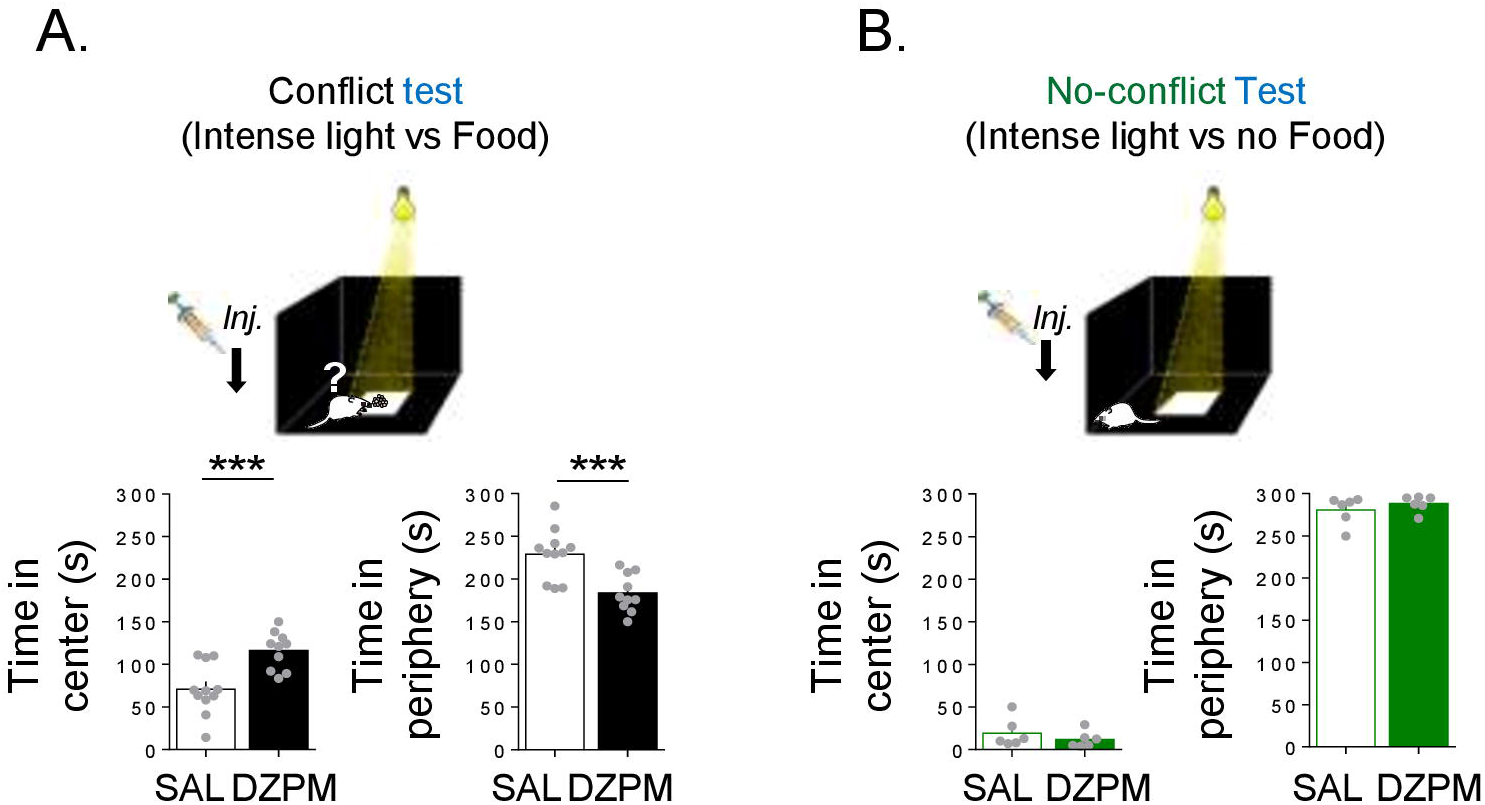
Diazepam increases time in center of an open field during conflict test without affecting no-conflict behavior. Rats placed in an open field arena were tested for innate exploratory behavior mediated either by conflict or no-conflict. ***A***, In the open field-mediated conflict test, diazepam-treated rats spent more time foraging for food (reward) in the brightly illuminated center of the arena (threat zone), and less time in the periphery (safe zone), as compared to the saline group (SAL, *n* = 11; DZMP, *n*= 10). ***B***, In the open field test where there is no food available (no reward) in the center of the arena (no-conflict), time in the brightly illuminated center and in the periphery were similar in saline solution and diazepam groups (SAL, n = 6; DZMP, *n* = 6). Error bars indicated SEM. *** *p* < 0.001.

## Discussion

We developed three conflict-mediated tasks for studying how rodents choose to confront threats to obtain rewards. We found that, regardless of whether competing cues where conditioned or innate, diazepam decreases active defensive responses triggered by threats, biasing choice towards reward-seeking behaviors during conflict. Thus, by using systemic pharmacological manipulations, we show that our choice-based tasks are valuable behavioral tools to study conflict behavior confronting threats to obtain rewards through active suppression of threat-related responses.

### Conflict-mediated tasks

Most previous conflict research has focused on cost-benefit decisions that involve choice between reward options (risky large rewards or small safe rewards) or reward punishment, thereby biasing behavior against risky reward-seeking behavior (Estes and Skinner 1941) and towards avoidance responses (Geller, Kulak et al. 1962, Jean-Richard-Dit-Bressel and McNally 2015, Orsini, Willis et al. 2016, Piantadosi, Yeates et al. 2017). In contrast, our three conflict-mediated tasks are set up to reward the act of overcoming threats, thereby biasing choice towards approach behaviors despite avoidance reactions. Our tasks involve rewarding risky behaviors based on training rats to distinguish between safe and risky locations to execute a choice behavior. Rats can choose between staying in the safe location to avoid risk or move to a risky location and obtain a reward. To obtain rewards they must overcome threats. One of our tasks involves extensive training to reward risky crossings (30 days to train rats to cross between “safe” zones in a straight alley despite threat), another task involves little training to reward risky step-down behavior (7 days to train rats to step-down from a safe platform to a threat zone) and a final task involves innate behaviors that reward risky foraging for food (one day of testing rats moving from safely exploring the periphery of an arena to feeding in a threat zone located in the center of an open field).

The crossing-mediated conflict task is a modified version of a task previously used to map self-stimulation brain sites. In a series of hallmark studies, Olds and Milner used such a task where the reward after crossing the grid was electrical stimulation (Olds and Milner 1954). Our task involves natural food reward rather than an electrical reward and the use of discrete learned cues to guide choice behavior. Our task requires that a hungry rat, previously trained to press a level for food cued by light, crosses an electrified grid to get to reach a lever on the other side of the straight alley that triggers a food dispenser. Our task allows for isolation of key variables in the same individual: appetitive drive indicated by latency to approach in safe no-conflict trials, defensive response indicated by freezing in the “threat” zone, and competition between aversive and appetitive drives indicated by latency to approach in conflict trials. Although using the reported training parameters, we have found (here and in previous pilot experiments) that most rats learn to cross despite cued threat (~80%), we have also encountered some rats that take too long or simply do not cross during conflict (~20%). Consequently, an additional advantage of this task is that it may allow identifying risk averse and risk approaching traits in individual rats. A final important distinction of our conflict-mediated tasks is that they allow contrasting choice behavior in conditions that involve conflict with those that occur in the absence of conflict.

Conflict tasks that contrast conflict against no-conflict conditions are scarce. Only two recently developed conflict tasks (to our knowledge) have focused on this important comparison. One task involves a single-trial test on a radial maze which evaluated rats that choose to either enter an arm in a maze that is associated with competing appetitive and aversive continuous contextual cues (conflict) or entering an arm that was not associated with any cued valences (neutral) (Nguyen, Schumacher et al. 2015). Unlike this task our crossing-mediated conflict test involves discriminating between discretely timed cues across several trials that involve conflict from those that do not, which may be useful to record the precise timing of changes in neuronal activity with respect to choice behavior (choice-point) where animals must commit to suppress defensive responses (or not) to obtain food. A second conflict task that contrasts conflict with no-conflict conditions focused on the expression of freezing defensive response during reward availability (conflict trials) compared to neutral trials (no-conflict) (Burgos-Robles, Kimchi et al. 2017). Unlike this task, our tasks focus on active (rather than passive) suppression of defensive responses to obtain rewards during conflict, which simulates real-life challenges more readily than other tasks (facing threats driven by foraging behavior). Also, contrasted with predator-based conflict models (Choi and Kim 2010, Walters, Jubran et al. 2019), the use of discrete conditioned signals in the crossing-mediated conflict task allows precise timing of choice behaviors triggered by threat and reward cues. Taken together, our three tasks allow for the separation of discrete variables controlling behavior in a drive competition setting in the same individual (separation of crossing behaviors that involve conflict and those that lack conflict) or in separate groups of animals (step-down avoidance memory and foraging behaviors mediated by conflict or lack of conflict). Thus, in the present study, the conflict against no-conflict comparison and use of discrete learned cues, allowed us to isolate the effects of experimental manipulations on conflict-mediated behavioral responses in the same individuals (conflict vs no-conflict trials in the same rats) and across groups of rats (conflict and no-conflict tests in separate groups of rats).

### Diazepam during conflict tests

Our finding that diazepam facilitates overcoming threats to obtain rewards, is consistent with the notion that this benzodiazepine reduces threat-related responses in conflict with reward-seeking behaviors. Previous conflict work has shown that diazepam increases: time spent in the open arms of elevated plus maze (Rex, Stephens et al. 1996, Chaouloff, Durand et al. 1997, Dalvi and Rodgers 1999), time spent in the center of an open field that includes food (Britton and Britton 1981, Bodnoff, Suranyi-Cadotte et al. 1989, Rex, Stephens et al. 1996), time spent in an illuminated but not a dark compartment (Chaouloff, Durand et al. 1997), foraging behavior (Walters, Jubran et al. 2019) as well as increased rates of punished reward responding (Vogel, Beer et al. 1971)(Paterson and Hanania 2010) and conditioned suppression during conflict (Kilts, Commissaris et al. 1981, Commissaris and Rech 1982). Consistently, we found that rats injected with diazepam decreased the time spent to cross and step-down from a safe zone to an electrified grid and increased the time spent in the center of a brightly lit open field that also includes food. Thus, previous results along with our present results using conflict tests, are consistent with the notion that diazepam has an anti-conflict effect (Liljequist and Engel 1984, Pericic and Pivac 1996, Rowlett, Lelas et al. 2006).

Although the anti-conflict effect has been interpreted also as anxiolytic effect (anxiety reducing) (Ljungberg, Lidfors et al. 1987, Beck and Fibiger 1995), we found that a low dose of diazepam distinctly affects choice behavior during conflict while leaving no-conflict behaviors intact (including anxiety-like behaviors, locomotion, motor coordination, lever pressing, feeding intake or passive freezing defensive responses). This diazepam effect exclusively on conflict contingencies, suggests that this drug reduces the probability of engaging predominant defensive behaviors only when conflict with reward stimuli is involved. An alternative explanation for the anti-conflict effect of diazepam is that this drug reduces the aversive signal (not necessarily involving anxiety) that emerges during conflict. Engaging in a situation that is simultaneously threatening and rewarding induces aversion. Consequently, conflict may represent an aversive signal (Dreisbach and Fischer 2012), which in some cases may induce anxiety. In our tasks, challenging rats to confront threats to obtain a reward may induce by itself an aversion signal that is reduced by diazepam. Thus, our findings support the idea that diazepam has an anti-conflict effect by reducing the conflict aversive signal and highlight the possibility of dissociating anti-conflict and antianxiety effects of diazepam.

### Using rewards to suppress fear

Confronting threats to obtain a reward involves suppressing defensive responses triggered by threats (fear) in pursuit of a goal. Prior work on fear suppression has focused on extinction, a passive process (Sotres-Bayon and Quirk 2010). However, the passive suppression that occurs in fear extinction is slow and temporary. Fear extinction can take hours to days to reach low (pre-conditioning) fear levels. Once fear responses have extinguished, fear can readily return (relapse) with the passage of time (spontaneous recovery), change of context (renewal) or a harmful reminder (reinstatement) (Bouton 2004). This has led to the investigation of alternative approaches to study regulation of fear (Bravo-Rivera and Sotres-Bayon 2020). One option is to study the active (rather than passive) suppression of fear. Novel behavioral paradigms in animals are required to understand the neurobiology of active fear suppression. In our three tasks, rats must quickly suppress their fear (conditioned or innate) to obtain a reward. Thus, our tasks represent a useful behavioral toolset to study how rats actively and rapidly suppress their fear in pursuit of rewards.

The neural circuits underlying the active suppression of fear (as in an “act of courage”) are barely known. Notably, a study in humans showed that the prefrontal and the temporal lobe (including the amygdala) are involved in voluntarily confronting fear of an approaching snake (Nili, Goldberg et al. 2010). In line with this, previous studies in humans suggest that the use of different voluntary cognitive coping strategies to inhibit fear (e.g. reappraisal) engage the same prefrontal-amygdala pathway and additional structures like the striatum and hippocampus (Hartley and Phelps 2010). However, due to the limitations of research in humans, these studies are not able to identify specific neural circuits and mechanisms involved in this type of fear suppression. Ongoing studies in our laboratory, using the tasks validated here, are beginning to reveal prefrontal and subcortical (including amygdala and striatum) brain circuits necessary to actively suppress fear responses when rewards are available during approach/avoidance conflict tests (Hernandez-Jaramillo and Sotres-Bayón 2018, Illescas-Huerta, Ramirez-Lugo et al. 2018). Thus, our validated conflict-mediated animal models may be valuable tools to study the intricate mechanisms of how the brain actively suppresses fear to obtain rewards and thereby may help find new approaches to treat mental disorders in humans characterized by deficits in decision-making in the face of conflicting emotional information.

## Author Contributions

Conceptualization, E.I-H, L.R-L, R.OS and F.S-B; Methodology, E.I-H, L.R-L, R.OS and F.S-B. Resources, J.Q and F.S-B; Supervision, F.S-B; Investigation, E.I-H, L.R-L and R.OS; Writing −Original Draft, E.I-H, L.R-L and F.S-B; Writing – Review & Editing, E.I-H, L.R-L, R.OS, J.Q and F.S-B; Funding Acquisition, F.S-B.

## Acknowledgments

We thank Dr. Christian Bravo-Rivera for comments on the manuscript and Sotres-Bayon laboratory members for helpful discussions.

## References

Amir, A., S. C. Lee, D. B. Headley, M. M. Herzallah and D. Pare (2015). “Amygdala Signaling during Foraging in a Hazardous Environment.” J Neurosci 35(38): 12994–13005.

Beck, C. H. and H. C. Fibiger (1995). “Conditioned fear-induced changes in behavior and in the expression of the immediate early gene c-fos: with and without diazepam pretreatment.” J Neurosci 15(1 Pt 2): 709–720.

Bodnoff, S. R., B. Suranyi-Cadotte, R. Quirion and M. J. Meaney (1989). “A comparison of the effects of diazepam versus several typical and atypical anti-depressant drugs in an animal model of anxiety.” Psychopharmacology (Berl) 97(2): 277–279.

Bouton, M. E. (2004). “Context and behavioral processes in extinction.” Learn Mem 11(5): 485–494.

Bravo-Rivera, C., C. Roman-Ortiz, E. Brignoni-Perez, F. Sotres-Bayon and G. J. Quirk (2014). “Neural structures mediating expression and extinction of platform-mediated avoidance.” J Neurosci 34(29): 9736–9742.

Bravo-Rivera, C. and F. Sotres-Bayon (2020). “From Isolated Emotional Memories to Their Competition During Conflict.” Front Behav Neurosci 14: 36.

Britton, D. R. and K. T. Britton (1981). “A sensitive open field measure of anxiolytic drug activity.” Pharmacol Biochem Behav 15(4): 577–582.

Burgos-Robles, A., E. Y. Kimchi, E. M. Izadmehr, M. J. Porzenheim, W. A. Ramos-Guasp, E. H. Nieh, A. C. Felix-Ortiz, P. Namburi, C. A. Leppla, K. N. Presbrey, K. K. Anandalingam, P. A. Pagan-Rivera, M. Anahtar, A. Beyeler and K. M. Tye (2017). “Amygdala inputs to prefrontal cortex guide behavior amid conflicting cues of reward and punishment.” Nat Neurosci 20(6): 824–835.

Calhoon, G. G. and K. M. Tye (2015). “Resolving the neural circuits of anxiety.” Nat Neurosci 18(10): 1394–1404.

Cardinal, R. N., J. A. Parkinson, J. Hall and B. J. Everitt (2002). “Emotion and motivation: the role of the amygdala, ventral striatum, and prefrontal cortex.” Neurosci Biobehav Rev 26(3): 321–352.

Chaouloff, F., M. Durand and P. Mormede (1997). “Anxiety- and activity-related effects of diazepam and chlordiazepoxide in the rat light/dark and dark/light tests.” Behav Brain Res 85(1): 27–35.

Choi, E. A., P. Jean-Richard-Dit-Bressel, C. W. G. Clifford and G. P. McNally (2019). “Paraventricular Thalamus Controls Behavior during Motivational Conflict.” J Neurosci 39(25): 4945–4958.

Choi, J. S. and J. J. Kim (2010). “Amygdala regulates risk of predation in rats foraging in a dynamic fear environment.” Proc Natl Acad Sci U S A 107(50): 21773–21777.

Commissaris, R. L. and R. H. Rech (1982). “Interactions of metergoline with diazepam, quipazine, and hallucinogenic drugs on a conflict behavior in the rat.” Psychopharmacology (Berl) 76(3): 282–285.

Dalvi, A. and R. J. Rodgers (1999). “Behavioral effects of diazepam in the murine plusmaze: flumazenil antagonism of enhanced head dipping but not the disinhibition of open-arm avoidance.” Pharmacol Biochem Behav 62(4): 727–734.

Dreisbach, G. and R. Fischer (2012). “Conflicts as aversive signals.” Brain Cogn 78(2): 94–98.

Estes, W. K. and B. F. Skinner (1941). “Some quantitative properties of anxiety.” Journal of Experimental Psychology 29(5): 390–400.

Friedman, A., D. Homma, L. G. Gibb, K. Amemori, S. J. Rubin, A. S. Hood, M. H. Riad and A. M. Graybiel (2015). “A Corticostriatal Path Targeting Striosomes Controls Decision-Making under Conflict.” Cell 161(6): 1320–1333.

Geller, I., J. T. Kulak, Jr. and J. Seifter (1962). “The effects of chlordiazepoxide and chlorpromazine on a punishment discrimination.” Psychopharmacologia 3: 374–385.

Goldstein, L. B. and J. N. Davis (1990). “Post-lesion practice and amphetamine-facilitated recovery of beam-walking in the rat.” Restor Neurol Neurosci 1(5): 311–314.

Hartley, C. A. and E. A. Phelps (2010). “Changing fear: the neurocircuitry of emotion regulation.” Neuropsychopharmacology 35(1): 136–146.

Hayden, B. Y. and M. E. Walton (2014). “Neuroscience of foraging.” Front Neurosci 8: 81.

Hernandez-Jaramillo, A. and F. Sotres-Bayón (2018). Basolateral amygdala, but not the orbitofrontal cortex, is necessary for motivational conflict responses guided by previous experiences. Society for Neuroscience. San Diego, CA.

Hu, H. (2016). “Reward and Aversion.” Annu Rev Neurosci 39: 297–324.

Illescas-Huerta, E., L. Ramirez-Lugo, R. Ordoñez-Sierra and F. Sotres-Bayon (2018). Prelimbic prefrontal cortex is necessary to face threats during a motivational conflict guided by learned, but not innate, stimuli. Society for Neuroscience. San Diego, CA.

Izquierdo, I., J. A. Quillfeldt, M. S. Zanatta, J. Quevedo, E. Schaeffer, P. K. Schmitz and J. H. Medina (1997). “Sequential role of hippocampus and amygdala, entorhinal cortex and parietal cortex in formation and retrieval of memory for inhibitory avoidance in rats.” Eur J Neurosci 9(4): 786–793.

Jean-Richard-Dit-Bressel, P. and G. P. McNally (2015). “The role of the basolateral amygdala in punishment.” Learn Mem 22(2): 128–137.

Johnson, D. N. (1978). “Effect of diazepam on food consumption in rats.” Psychopharmacology (Berl) 56(1): 111–112.

Kilts, C. D., R. L. Commissaris and R. H. Rech (1981). “Comparison of anti-conflict drug effects in three experimental animal models of anxiety.” Psychopharmacology (Berl) 74(3): 290–296.

LeDoux, J. E. (2000). “Emotion circuits in the brain.” Annu Rev Neurosci 23: 155–184.

Liljequist, S. and J. A. Engel (1984). “The effects of GABA and benzodiazepine receptor antagonists on the anti-conflict actions of diazepam or ethanol.” Pharmacol Biochem Behav 21(4): 521–525.

Ljungberg, T., L. Lidfors, M. Enquist and U. Ungerstedt (1987). “Impairment of decision making in rats by diazepam: implications for the “anticonflict” effects of benzodiazepines.” Psychopharmacology (Berl) 92(4): 416–423.

Miller, S. M., D. Marcotulli, A. Shen and L. S. Zweifel (2019). “Divergent medial amygdala projections regulate approach-avoidance conflict behavior.” Nat Neurosci 22(4): 565–575.

Mobbs, D., P. C. Trimmer, D. T. Blumstein and P. Dayan (2018). “Foraging for foundations in decision neuroscience: insights from ethology.” Nat Rev Neurosci 19(7): 419–427.

Moscarello, J. M. and J. E. LeDoux (2013). “Active avoidance learning requires prefrontal suppression of amygdala-mediated defensive reactions.” J Neurosci 33(9): 3815–3823.

Naruse, T. and R. Ishii (1995). “Relationship between histamine receptors in the brain and diazepam-induced hyperphagia in rats.” Pharmacol Biochem Behav 51(4): 923–927.

Nguyen, D., A. Schumacher, S. Erb and R. Ito (2015). “Aberrant approach-avoidance conflict resolution following repeated cocaine pre-exposure.” Psychopharmacology (Berl) 232(19): 3573–3583.

Nili, U., H. Goldberg, A. Weizman and Y. Dudai (2010). “Fear thou not: activity of frontal and temporal circuits in moments of real-life courage.” Neuron 66(6): 949–962.

Olds, J. and P. Milner (1954). “Positive reinforcement produced by electrical stimulation of septal area and other regions of rat brain.” J Comp Physiol Psychol 47(6): 419–427.

Orsini, C. A., M. L. Willis, R. J. Gilbert, J. L. Bizon and B. Setlow (2016). “Sex differences in a rat model of risky decision making.” Behav Neurosci 130(1): 50–61.

Paterson, N. E. and T. Hanania (2010). “The modified Geller-Seifter test in rats was insensitive to GABAB receptor positive modulation or blockade, or 5-HT1A receptor activation.” Behav Brain Res 208(1): 258–264.

Pericic, D. and N. Pivac (1996). “Effects of diazepam on conflict behaviour and on plasma corticosterone levels in male and female rats.” Naunyn Schmiedebergs Arch Pharmacol 353(4): 369–376.

Piantadosi, P. T., D. C. M. Yeates, M. Wilkins and S. B. Floresco (2017). “Contributions of basolateral amygdala and nucleus accumbens subregions to mediating motivational conflict during punished reward-seeking.” Neurobiol Learn Mem 140: 92–105.

Prut, L. and C. Belzung (2003). “The open field as a paradigm to measure the effects of drugs on anxiety-like behaviors: a review.” Eur J Pharmacol 463(1-3): 3–33.

Rangel, A., C. Camerer and P. R. Montague (2008). “A framework for studying the neurobiology of value-based decision making.” Nat Rev Neurosci 9(7): 545–556.

Rex, A., D. N. Stephens and H. Fink (1996). ““Anxiolytic” action of diazepam and abecarnil in a modified open field test.” Pharmacol Biochem Behav 53(4): 1005–1011.

Rodgers, R. J., B. J. Cao, A. Dalvi and A. Holmes (1997). “Animal models of anxiety: an ethological perspective.” Braz J Med Biol Res 30(3): 289–304.

Rowlett, J. K., S. Lelas, W. Tornatzky and S. C. Licata (2006). “Anti-conflict effects of benzodiazepines in rhesus monkeys: relationship with therapeutic doses in humans and role of GABAA receptors.” Psychopharmacology (Berl) 184(2): 201–211.

Schmitt, U. and C. Hiemke (1998). “Strain differences in open-field and elevated plus-maze behavior of rats without and with pretest handling.” Pharmacol Biochem Behav 59(4): 807–811.

Schumacher, A., F. R. Villaruel, A. Ussling, S. Riaz, A. C. H. Lee and R. Ito (2018). “Ventral Hippocampal CA1 and CA3 Differentially Mediate Learned Approach-Avoidance Conflict Processing.” Curr Biol 28(8): 1318–1324 e1314.

Sotres-Bayon, F. and G. J. Quirk (2010). “Prefrontal control of fear: more than just extinction.” Curr Opin Neurobiol 20(2): 231–235.

Stefanski, R., W. Palejko, W. Kostowski and A. Plaznik (1992). “The comparison of benzodiazepine derivatives and serotonergic agonists and antagonists in two animal models of anxiety.” Neuropharmacology 31(12): 1251–1258.

Tye, K. M. (2018). “Neural Circuit Motifs in Valence Processing.” Neuron 100(2): 436–452.

Verharen, J. P. H., M. W. van den Heuvel, M. Luijendijk, L. Vanderschuren and R. A. H. Adan (2019). “Corticolimbic Mechanisms of Behavioral Inhibition under Threat of Punishment.” J Neurosci 39(22): 4353–4364.

Vogel, J. R., B. Beer and D. E. Clody (1971). “A simple and reliable conflict procedure for testing anti-anxiety agents.” Psychopharmacologia 21(1): 1–7.

Walsh, R. N. and R. A. Cummins (1976). “The Open-Field Test: a critical review.” Psychol Bull 83(3): 482–504.

Walters, C. J., J. Jubran, A. Sheehan, M. T. Erickson and A. D. Redish (2019). “Avoid-approach conflict behaviors differentially affected by anxiolytics: implications for a computational model of risky decision-making.” Psychopharmacology (Berl) 236(8): 2513–2525.

